# The effect of enriched versus depleted housing on eucalyptus smoke-induced cardiovascular dysfunction in mice

**DOI:** 10.1101/2024.02.26.582161

**Authors:** Molly E. Harmon, Michelle Fiamingo, Sydnie Toler, Kaleb Lee, Yongho Kim, Brandi Martin, Ian Gilmour, Aimen K. Farraj, Mehdi S. Hazari

**Author notes:** Corresponding author: Mehdi S. Hazari,109 TW Alexander Blvd, B-105 Research Triangle Park, NC 27711. Disclaimer: This paper has been reviewed and approved for release by the Center for Public Health and Environmental Assessment, U.S. Environmental Protection Agency. Approval does not signify that the contents necessarily reflect the views and policies of the U.S. EPA, nor does mention of trade names. This version of this paper has not yet been through peer-review. The authors report there are no competing interests to declare. All data are available on EPA ScienceHub.

## Abstract

Living conditions play a major role in health and well-being, particularly for the cardiovascular and pulmonary systems. Depleted housing contributes to impairment and development of disease, but how it impacts body resiliency during exposure to environmental stressors is unknown. This study examined the effect of depleted (DH) versus enriched housing (EH) on cardiopulmonary function and subsequent responses to wildfire smoke. Two cohorts of healthy female mice, one of them surgically implanted with radiotelemeters for the measurement of electrocardiogram, body temperature (Tco) and activity, were housed in either DH or EH for 7 weeks. Telemetered mice were exposed for 1 hour to filtered air (FA) and then flaming eucalyptus wildfire smoke (WS) while untelemetered mice, which were used for ventilatory assessment and tissue collection, were exposed to either FA or WS. Animals were continuously monitored for 5-7 days after exposure. EH prevented a decrease in Tco after radiotelemetry surgery. EH mice also had significantly higher activity levels and lower heart rate during and after FA and WS. Moreover, EH caused a decreased number of cardiac arrhythmias during WS. WS caused ventilatory depression in DH mice but not EH mice. Housing enrichment also upregulated the expression of cardioprotective genes in the heart. The results of this study indicate that housing conditions impact overall health and cardiopulmonary function. More importantly, depleted housing appears to worsen the response to air pollution. Thus, non-chemical factors should be considered when assessing the susceptibility of populations, especially when it comes to extreme environmental events.

## Introduction

The overall health of the human body and its resistance to adverse responses, whether due to general stress or environmental exposures, is determined by numerous daily and at times persistent modifiers that alter susceptibility and risk over the long term, particularly for cardiovascular disease (Joseph et al. 2020). In recent years, the prominent role of social factors in health has become increasingly recognized (Hacker et al. 2022). These social determinants of health include, among other things, access to education and healthcare, economic stability, and living conditions. Poor housing environment has drawn particular attention because it causes heightened risk of disease in certain vulnerable and underserved populations, (Krieger and Higgins 2002).

In 2020, the American Heart Association (AHA) cited an association between housing status and cardiovascular health, while simultaneously emphasizing that knowledge gaps for how certain housing conditions affect cardiovascular health remain (Sims et al. 2020). It has already been shown that the living environment contributes to the progression of hypertension (Ford et al. 2016) and coronary artery disease (Diez Roux et al. 2001) and worsens co-morbidities (Ganatra et al. 2022). These deleterious effects are likely caused by increased psychosocial stress and subsequent underlying changes in the regulation of the cardiovascular system. Epidemiological data show that poor housing conditions early in life result in higher-than-normal cortisol levels (Blair et al. 2011) suggesting subsequent responsiveness to stressors such as environmental pollutants could be worsened.

We have previously characterized the adverse cardiopulmonary impacts of wildfire smoke (Kim et al. 2014; Thompson et al. 2018; McGee-Hargrove et al. 2019; Martin et al. 2020), which has become a persistent problem over the last few years. Moreover, the health effects of wildfire smoke are underscored by the fact that certain populations are disproportionately harmed due to socio-economic factors (Rappold et al. 2012), which once again, points to the role of certain factors like living conditions as important determinants of body resiliency. Unfortunately, data on the direct impact of living conditions on cardiovascular responses to environmental exposures is still inadequate, especially if the goal is to establish biological plausibility and develop subsequent mitigation strategies.

Housing quality and social environment have already been identified as major contributors to cardiovascular health, and although their effect has been documented in rodents (Schmidt et al. 2022; Watanabe et al. 2022), there are no studies examining how they alter responsiveness to an environmental stressor. For instance, it has already been shown that housing rats in metabolic rather than standard cages result in mild and reversible cardiovascular changes (Giral et al. 2022). In addition, a previous study showed that enriched housing improved the behavior and overall health of wildtype mice (Kulesskaya et al. 2011). The purpose of this current study was to determine whether housing conditions affect the response of mice to a single wildfire-related smoke exposure. We hypothesized that enriched housing would alter the baseline cardiovascular function of mice and reduce the impact of smoke exposure when compared to mice housed in a depleted environment.

## Materials and Methods

### Animals

Eight-week-old female C57BL/6 mice (body weight = 18.8 ± 0.2 g) were used in this study (Jackson Laboratory – Bar Harbor, ME). Mice were initially housed five per cage and maintained on a 12-hr light/dark cycle at approximately 22°C and 50% relative humidity in an AAALAC–approved facility. Food (Prolab RMH 3000; PMI Nutrition International, St. Louis, MO) and water were provided ad libitum. All protocols were approved by the Institutional Animal Care and Use Committee of the U.S. Environmental Protection Agency and are in accordance with the National Institutes of Health Guides for the Care and Use of Laboratory Animals. The animals were treated humanely and with regard for alleviation of suffering.

### Experimental groups and housing conditions

Mice were randomly assigned to one of two groups: (1) depleted/unenriched (DH) or (2) enriched (EH) housing. All animals were housed five per cage with alpha-dri bedding and a single nestlet (i.e., standard – housing) (Lab Supply, Durham, NC) for one week upon arrival to the facility and thereafter separated and moved to either DH or EH for 7 weeks. A subset of animals (Cohort A) in each of the housing groups underwent surgical implantation of radiotelemeters for the measurement of activity, core body temperature (Tco) and electrocardiogram (ECG) during week 3 of the housing regimen (n = 11-12), while the rest (Cohort B) remained untelemetered for the measurement of ventilatory function and tissue collection at necropsy (n = 8-12). DH mice implanted with radiotelemeters were housed individually in a cage with only bedding whereas radiotelemetered mice in the EH group were housed individually in a cage with bedding, a nestlet, hut, running saucer, tunnel and nylabone (chew toy) to mimic positive human psychosocial living conditions (greenspace, physical enrichment, etc). Although mice in our facility are normally housed five per cage for socialization purposes, telemetered animals must be housed individually to prevent any complications in their post-surgical health and/or interference in telemetry signal. Untelemetered DH mice were housed five per cage with only bedding while untelemetered EH mice were housed five per cage with bedding, two nestlets, a hut, running saucer, tunnel and two nylabones. All the animals were maintained in their respective housing conditions for 7 weeks (Figure 1 – ‘Week 1’ marks the start of the housing regimen). All Cohort A animals were exposed to filtered air (FA) first (i.e. air sham) and eucalyptus wildfire smoke (WS) thereafter (separated by four days), and thus, served as their own controls. Cohort B animals underwent ventilatory function assessments after an initial week 5 exposure to FA (air sham) and then WS (i.e. served as their own controls, FA → WS) during week 6-7, and were then necropsied 24 hours later. A parallel set of animals in Cohort B were only exposed to FA in week 6-7 (FA → FA) and necropsied thereafter.

**Figure 1.**
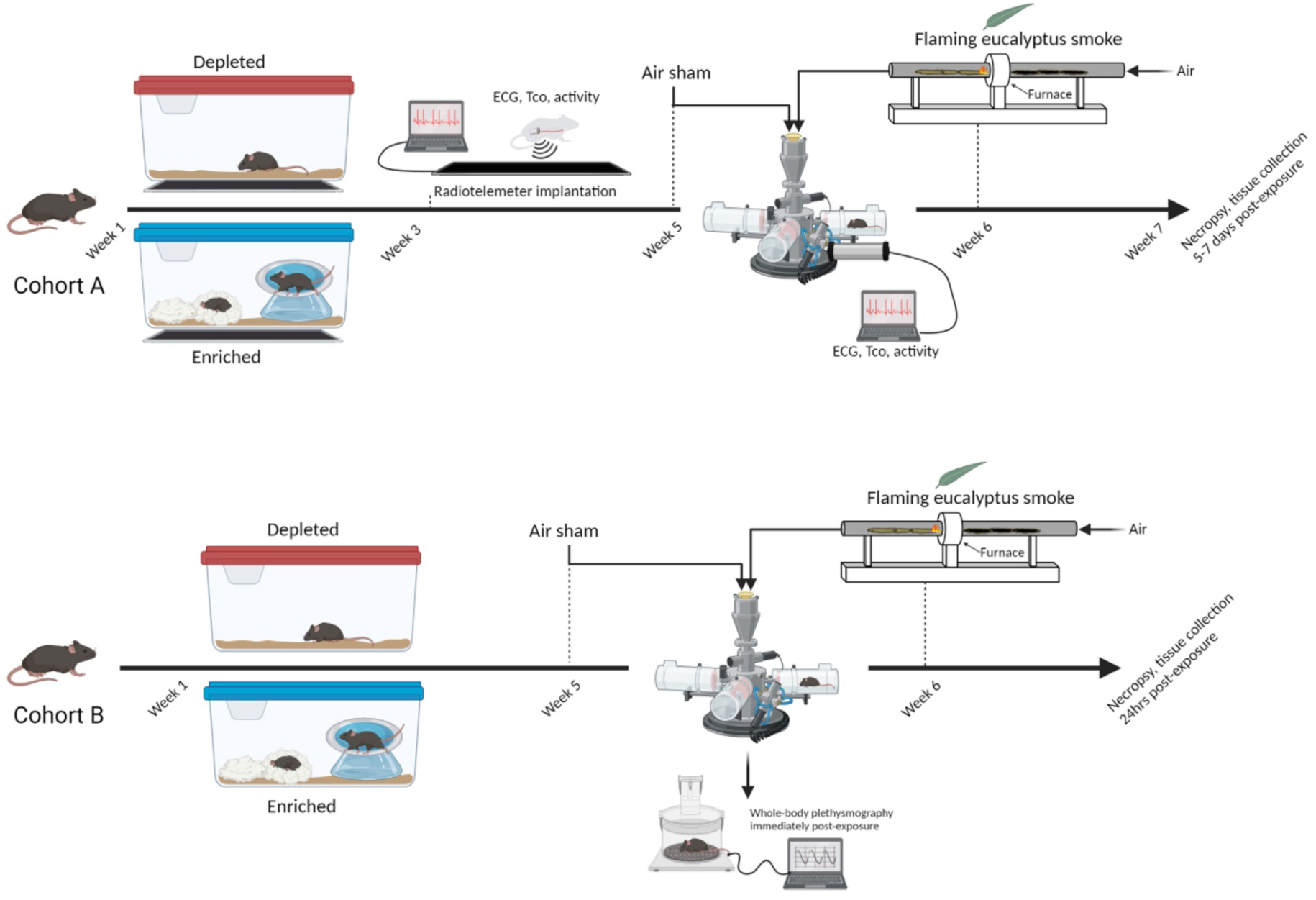
Study design and schedule. Mice were assigned to either depleted/unenriched (DH) or enriched (EH) housing. A subset of animals (Cohort A) in each of the housing groups underwent surgical implantation of radiotelemeters during week 3, while the rest (Cohort B) remained untelemetered. All the animals were maintained in their respective housing conditions for approximately 6-7 weeks. All Cohort A animals were exposed to filtered air (FA) during week 5 and eucalyptus wildfire smoke (WS) during week 6, and then monitored for 5-7 days and then necropsied. Cohort B animals underwent ventilatory function assessments after an initial week 5 exposure to FA and then WS during week 6 and were then necropsied 24 hours later. A parallel set of animals in Cohort B were only exposed to FA in week 6 and necropsied thereafter. (Image created in Biorender)

### Surgical implantation of radiotelemeters

Animals were weighed and then anesthetized using inhaled isoflurane (Isothesia, Butler Animal Health Supply, Dublin, OH). Anesthesia was induced by spontaneous breathing of 2.5% isoflurane in pure oxygen at a flow rate of 1 L/min and then maintained by 1.5% isoflurane in pure oxygen at a flow rate of 0.5 L/min; all animals received the analgesic buprenorphrine (0.03 mg/kg, i.p. Henry Schein, Melville, NY). Briefly, using aseptic technique, each animal was implanted subcutaneously with a radiotelemeter (ETA-F10, Data Sciences International, St Paul, MN); the transmitter was placed under the skin to the right of the midline on the dorsal side. The two electrode leads were then tunneled subcutaneously across the lateral dorsal sides; the distal portions were fixed in positions that approximated those of the lead II of a standard ECG. Body heat was maintained both during and immediately after the surgery. Animals were given food and water post-surgery and were housed individually. All animals were allowed 7-10 days to recover from the surgery and reestablish circadian rhythms.

### Radiotelemetry data acquisition

Radiotelemetry methodology (Data Sciences International, Inc., St. Paul, MN) was used to track changes in activity, which is unitless and measures animal movement, core body temperature (Tco) and cardiovascular function by monitoring heart rate (HR) and ECG waveforms immediately following telemeter implantation, during the 3-4 weeks of housing regimen, through air sham and WS exposure, and 5-7 days after WS. This methodology provided continuous monitoring and collection of physiologic data from individual mice to a remote receiver. Sixty-second ECG segments were recorded every 5 minutes during the pre– and post-exposure periods. ECG was recorded continuously during exposure (air sham, baseline and wildfire smoke – see below); HR was automatically obtained from the waveforms (Dataquest ART Software, version 3.01, Data Sciences International, St. Paul, MN, USA).

### Electrocardiogram (ECG) analysis

Ponemah software (Data Sciences International, St. Paul, MN) was used to visualize individual ECG waveforms, analyze, and quantify ECG segment durations and areas, as well as identify cardiac arrhythmias. Briefly, P-wave, QRS complex, and T-wave were identified for individual ECG waveforms. Proper identification of ECG waveform by the software and positioning of landmarks was validated and artifacts were removed. All ECG streams with less than 10 s of identifiable cardiac cycles were excluded from ECG parameter calculations.

### Heart rate variability analysis

Heart rate variability (HRV) was calculated as the mean of the differences between sequential RRs for the complete set of ECG waveforms using Ponemah. After removing artifacts and signal segments that were not easily identifiable, mean time between successive QRS complex peaks (RR interval), mean HR, and mean HRV-analysis–generated time-domain measures were acquired for each 1-min stream of ECG waveforms. The time-domain measures included standard deviation of the time between normal-to-normal beats (SDNN), and root mean squared of successive differences (RMSSD). SDNN represents overall HRV, while RMSSD represents parasympathetic influence over HRV. HRV analysis was also conducted in the frequency domain using a fast-Fourier transform. For frequency domain analysis, the signal was analyzed with a Hamming window for segment lengths of 512 samples with 50% overlapping. The spectral power obtained from this transformation represents the total harmonic variability for the frequency range being analyzed. In this study, the spectrum was divided into low-frequency (LF) and high-frequency (HF) regions with frequency bands assigned as 0.15-1.5 and 1.5-5 Hz respectively. LF is generally believed to represent a combination of sympathetic and parasympathetic activity, while HF indicates cardiac parasympathetic (vagal) activity (Ori 1992). All ECG streams with less than 1 min of identifiable RR intervals were excluded from HRV analysis. Additionally, thorough visual inspection was conducted to identify and exclude arrhythmias and artifacts.

### Whole-body plethysmography

Ventilatory function was assessed in awake, unrestrained mice in Cohort B using a whole-body plethysmograph (Data Sciences International, St. Paul, MN). Assessments were performed immediately after air sham exposure and immediately following exposure to WS. The plethysmograph pressure was monitored using Biosystems XA software (Buxco Electronics Inc., Wilmington, NC). Using respiratory-induced fluctuations in ambient pressure, ventilatory parameters including tidal volume (TV), breathing frequency (f), inspiratory time (Ti), expiratory time (Te), minute volume (MV) and enhanced pause (Penh), which is a measure of ventilatory timing and can indicate airway irritation, were calculated, and recorded on a breath-by-breath basis. Changes from post-FA to post-WS were determined for each animal.

### Tube furnace exposure system, eucalyptus wildfire smoke, particulate matter (PM) and gas monitoring and sampling

All animals were acclimated twice in thirty-minute increments to a whole-body inhalation tube at room temperature over a 2-day period leading up to exposure. A one-hour air sham exposure was conducted for each animal in Cohorts A and B. We used eucalyptus fuel which was purchased as writing pen blanks (rectangles at 0.75 inches square by 6 inches long) (Woodworkers source Arizona). This wood was processed through a gasoline powered wood shredder (Echo bearcat model number SC3206). The shredder was cleaned between uses so no cross contamination would occur.

Animals were exposed once to WS generated using an automated control tube furnace system (Kim et al. 2018). The inhalation exposures were conducted with mice being individually housed in larger rat restraint tubes that were inserted into ports of a “nose and mouth only” exposure chamber, allowing unrestrained free movement. Mice were exposed once to the biomass smoke (flaming) for one hour. We monitored carbon dioxide (CO_2_) and carbon monoxide (CO) levels using a non-dispersive infrared analyzer (Model: 602 CO/CO_2_; CAI Inc., Orange, CA) and nitrogen oxides (NOx) using a chemiluminescent analyzer (Model: 42i NO/NO_2_/NOx; Thermo Scientific, Franklin, MA). Because CO in biomass smoke can elicit its own health impacts in terms of altering respiratory rate, vascular function and other endpoints, we elected to hold the exposures to concentrations of CO below 60 ppm. Preliminary testing during combustion runs showed PM concentrations of approximately 4 mg/m^3^. We also collected PM on a glass-fiber filter installed in an exhaust line of the inhalation chamber to determine mean PM concentrations gravimetrically by weighing the filter before and after inhalation exposure. The real-time measurements of WS properties and engineering parameters (e.g., temperature, RH, static pressure, and flow rate) were continuously monitored, recorded, and displayed using data acquisition software (Dasylab version 13.0, National Instruments, Austin, TX)

### RNA Isolation and Expression Analysis

Total RNA was isolated from cardiac tissue using Trizol reagent (Invitrogen, Waltham, MA) following a phenol-chloroform extraction protocol. RNA concentrations and integrity were confirmed using a NanoDrop spectrophotometer (Thermo Scientific, Wilmington, DE) and Qubit fluorometer BR (Broad Range) assay kit (Thermofisher) prior to shipment to Nanostring for gene expression quantification. For gene expression analysis, we used a customized NanoString’s nCounter® Elements TagSet panel of 100 genes coding for proteins related to cardiac contraction, ion signaling, oxidative stress, hemodynamic control and other cardiac function. This system is based on direct multiplexed quantification of nucleic acids enabling the profiling of hundreds of unique target mRNAs in a single reaction. Gene expression was quantified using nSolver Analysis Software, an integrated analysis platform for storage, custom QC, and custom normalization of nCounter data. Data was analyzed with a HyperScale architecture by ROSALIND, Inc. (San Diego, CA). Read Distribution percentages, violin plots, identity heatmaps, and sample MDS plots were generated as part of the QC step. Normalization, fold changes and p-values were calculated using criteria provided by Nanostring. ROSALIND® follows the nCounter® Advanced Analysis protocol of dividing counts within a lane by the geometric mean of the normalizer probes from the same lane. Housekeeping probes used for normalization (i.e., actin beta (ACTB), beta-2-microglobulin (B2M), and glyceraldehyde-3-phosphate dehydrogenase (GAPDH)) were selected based on the geNorm algorithm as implemented in the NormqPCR R library1. Data represented the mean of normalized counts and were expressed as fold-change. We selected genes that were lower than 1.25-fold and higher than 1.25-fold and also examined the changes at +/− 1.15-fold. DH versus EH differences in expression levels were considered significantly different if the fold changes yielded a p < 0.05, corrected for multiple comparisons. Gene ontology (GO)-based pathway analyses were also performed.

### Tissue Collection and Analysis

Mice were deeply anesthetized with an intra-peritoneal injection of a sodium pentobarbital/phenytoin solution and euthanized 24 hours after exposure and blood and bronchoalveolar lavage fluid (BAL) were collected, processed, and analyzed as previously described (Kurhanewicz et al. 2018). Multiple biochemical markers were assessed in the serum and BAL. To assess cardiopulmonary inflammation, injury and oxidative stress, multiple biochemical markers and inflammatory cells were assayed. Lavage supernatants were analyzed for lactate dehydrogenase (LDH), N-acetyl-b-d-glucosaminidase (NAG), superoxide dismutase (SOD), and total protein. Serum supernatants were analyzed for cholesterol, creatine kinase (CK), C-reactive protein (CRP), glutathione peroxidase, glutathione reductase, high-density lipoprotein, and LDH.

### Statistics

All data are expressed as means ± SEM. Statistical analyses were performed using Sigmaplot 13.0 (Systat Software, San Jose, CA) software. Sample size analysis was based upon the (*k*) number of experimental groups, a significance level = 0.05, a power = 0.8, and the effect size index (*f*), which is derived by multiplying the expected effect size (d) by the standard deviation (SD). Using these, an “*n*” of 8 mice per group was determined, but more included in the event animals were lost. We performed tests of normality (Shapiro-Wilk) for all continuous variables and used parametric methods of analysis where appropriate. A linear mixed model with least squares means and repeated measures ANOVA was used to determine which interactions (i.e., time x housing, housing x exposure) were statistically significant. Multiple comparison adjustment for the p values and confidence limits for the differences between the least squares means was done using a *post hoc* test (Tukey’s). The statistical significance was set at P < 0.05.

## Results

### Exposure characteristics

Table 1 shows the characteristics of the flaming eucalyptus wildfire smoke exposures. The mean particulate matter (PM_2.5_) concentration was 3.3 ± 0.2 mg/m^3^, which was comparable to the level used by our colleagues in previous mouse studies (Hargrove et al. 2019) and slightly less than the threshold used by our colleagues to investigate cardiopulmonary complications in humans (DeFlorio-Barker et al. 2019). Levels of CO, CO_2_, and NOx were 87.0 ± 4.9 ppm, 3040 ± 56.8 ppm and 2.4 ± 0.1 ppm, respectively. Modified combustion efficiency (MCE), defined as MCE (%) = (ΔCO2 / (ΔCO2 + ΔCO)) × 100, where ΔCO_2_ and ΔCO are the excess concentrations of CO_2_ and CO, was calculated to confirm that the flaming exposure conditions (>95%) were achieved. Other component levels were comparable to our previous flaming eucalyptus exposures (Hargrove et al. 2019).

**Table 1.**
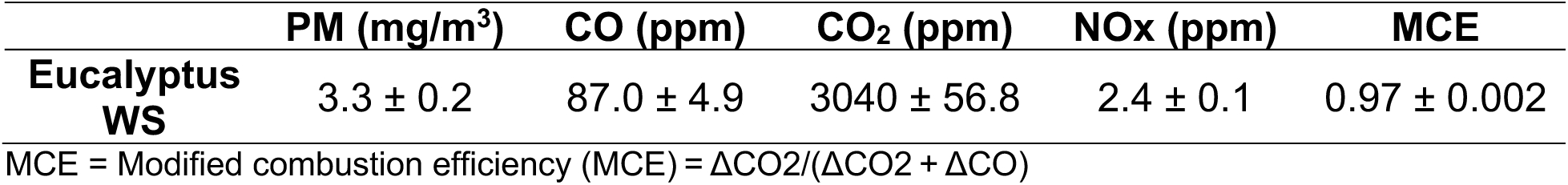
Exposure details.

|  | PM (mg/m<sup>3</sup>) | CO (ppm) | CO<sub>2</sub> (ppm) | NOx (ppm) | MCE |
| --- | --- | --- | --- | --- | --- |
| **Eucalyptus**<br>**WS** | 3.3 ± 0.2 | 87.0 ± 4.9 | 3040 ± 56.8 | 2.4 ± 0.1 | 0.97 ± 0.002 |

MCE = Modified combustion efficiency (MCE) = $\Delta\text{CO}_2 / (\Delta\text{CO}_2 + \Delta\text{CO})$

### Body weight

There was no difference in body weights at Week 1 when animals were separated into DH (20.8 ± 0.3 g) and EH (21.0 ± 0.2g). They were measured again for Cohort A animals at telemetry surgery. All the animals, Cohorts A (Fig. 2A) and B (data not shown), experienced an increase in body weight during the course of the study. In Cohort A, EH mice had less weight gain when compared to DH after radiotelemetry surgery (Fig. 2B). There was no difference in weights or change in weight between DH and EH mice in Cohort B.

**Figure 2.**
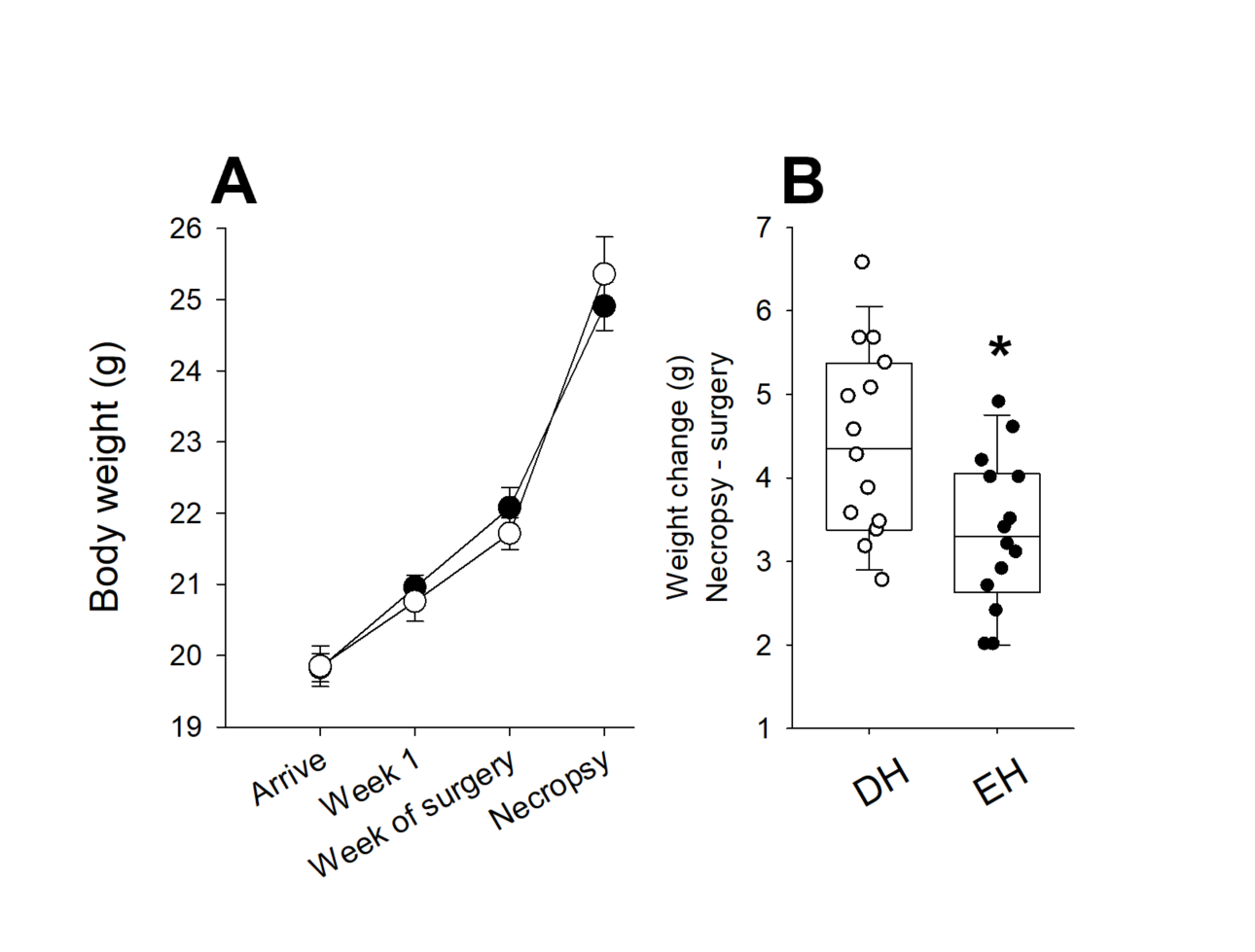
Living in a depleted housing environment impacts change in body weight. A. There was no difference in body weights at Week 1 when animals were separated into DH (open circles) and EH (filled circles). All the animals, Cohorts A mice experienced an increase in body weight during the course of the study. B. In Cohort A, EH mice had less weight gain when compared to DH after radiotelemetry surgery. n = 18-20; * Significantly different from DH, p < 0.05.

### Core body temperature

Body temperature was maintained during radiotelemeter surgery and monitored continuously thereafter. EH mice had significantly higher Tco (i.e., Tco dropped significantly in DH mice post-surgery), which returned to normal more quickly immediately after surgery when compared to DH (Figure 3A). Both groups had a nighttime increase and daytime decrease in Tco, which coincided with circadian rhythm. There was no difference in the Tco between the groups after that (data not shown), although DH animals had an increasing trend (p = 0.08) during air sham (Figure 3B).

**Figure 3.**
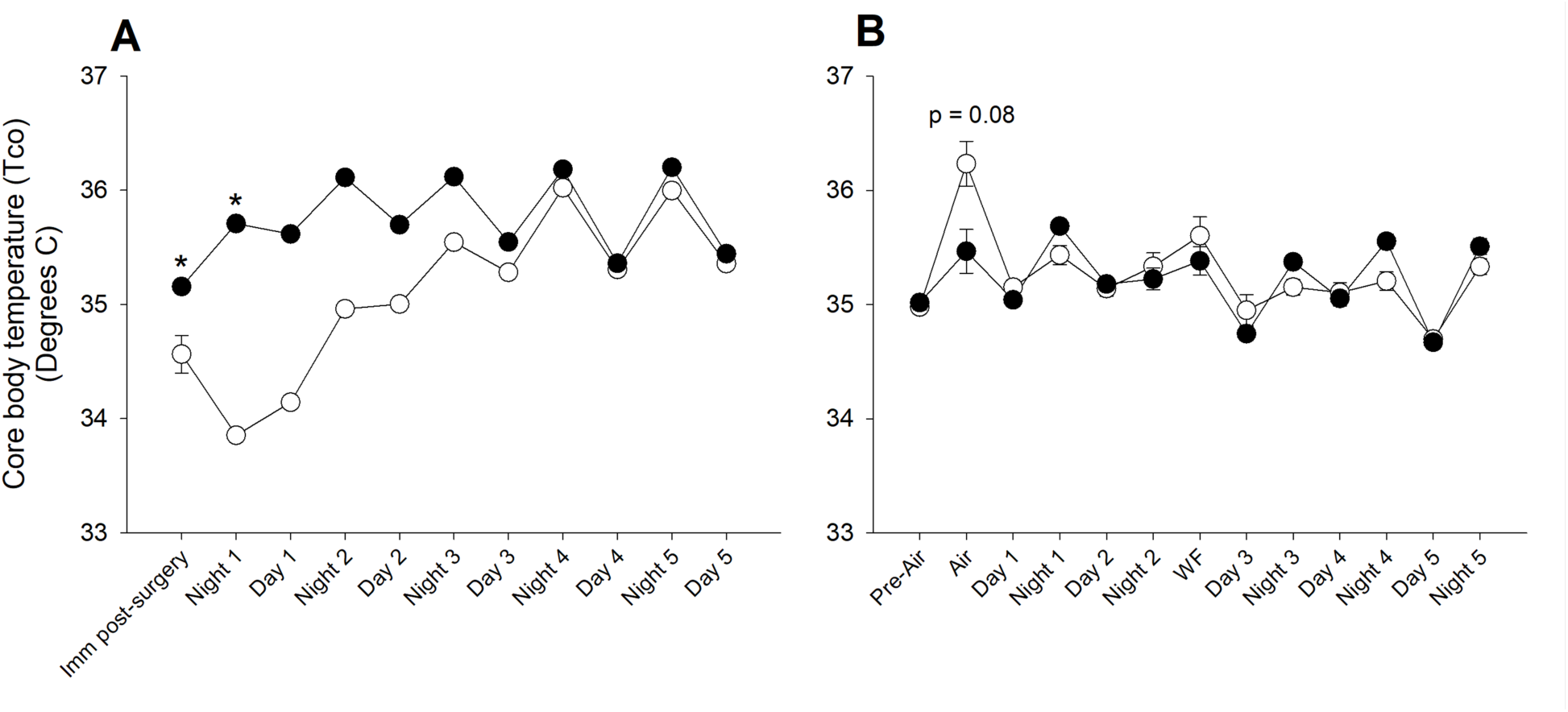
Housing conditions affect core body temperature. A. Core body temperature was normal in Cohort A mice housed in EH (filled circles) after radiotelemetry surgery when compared to DH (open circles), which had a significant decrease. B. There were no core body temperature differences between DH and EH after air sham or WS exposure, however, there was an increasing trend in the former. n = 10-12; * Significantly different from DH, p < 0.05.

### Heart rate and cardiac arrhythmia

Radiotelemetry data acquisition, which included HR, began post-surgery; specific relevant segments are shown in Figure 4. In general, the impact of housing conditions on HR did not depend on time/period. DH mice had significantly lower HR in the nighttime immediately following surgery. Although both EH and DH mice had fluctuating HR until three days post-surgery, they re-established circadian pattern (higher HR at night and lower during the day) thereafter. HR fluctuated again before the air sham (Pre-sham) exposure, likely due to acclimation in the chamber; however, EH animals had lower HR than DH in general. During the early part of air-sham exposure, EH mice had significantly lower HR when compared to DH. Although HR was similar for both groups by the end of the air sham, DH animals had a significant increase immediately thereafter (Post-sham); this persisted for approximately two days. Interestingly, day and nighttime HR before WS exposure (Pre-WS) was lower in EH animals than DH. During WS exposure, HR steadily decreased for both groups; however, similar to the air sham, DH animals experienced a significant increase at the end and immediately after (Post-WS). HR remained elevated in DH animals during the daytime four to five days after WS. Mice living in the enriched housing had less change in their HR at the end of the one-hour WS exposure relative to their air sham when compared to those in depleted housing (Figure 4B). During exposure, DH mice also had a significantly greater number of non-conducted p-waves, which are blocks in normal cardiac electrical conduction (Figure 4C and D).

**Figure 4.**
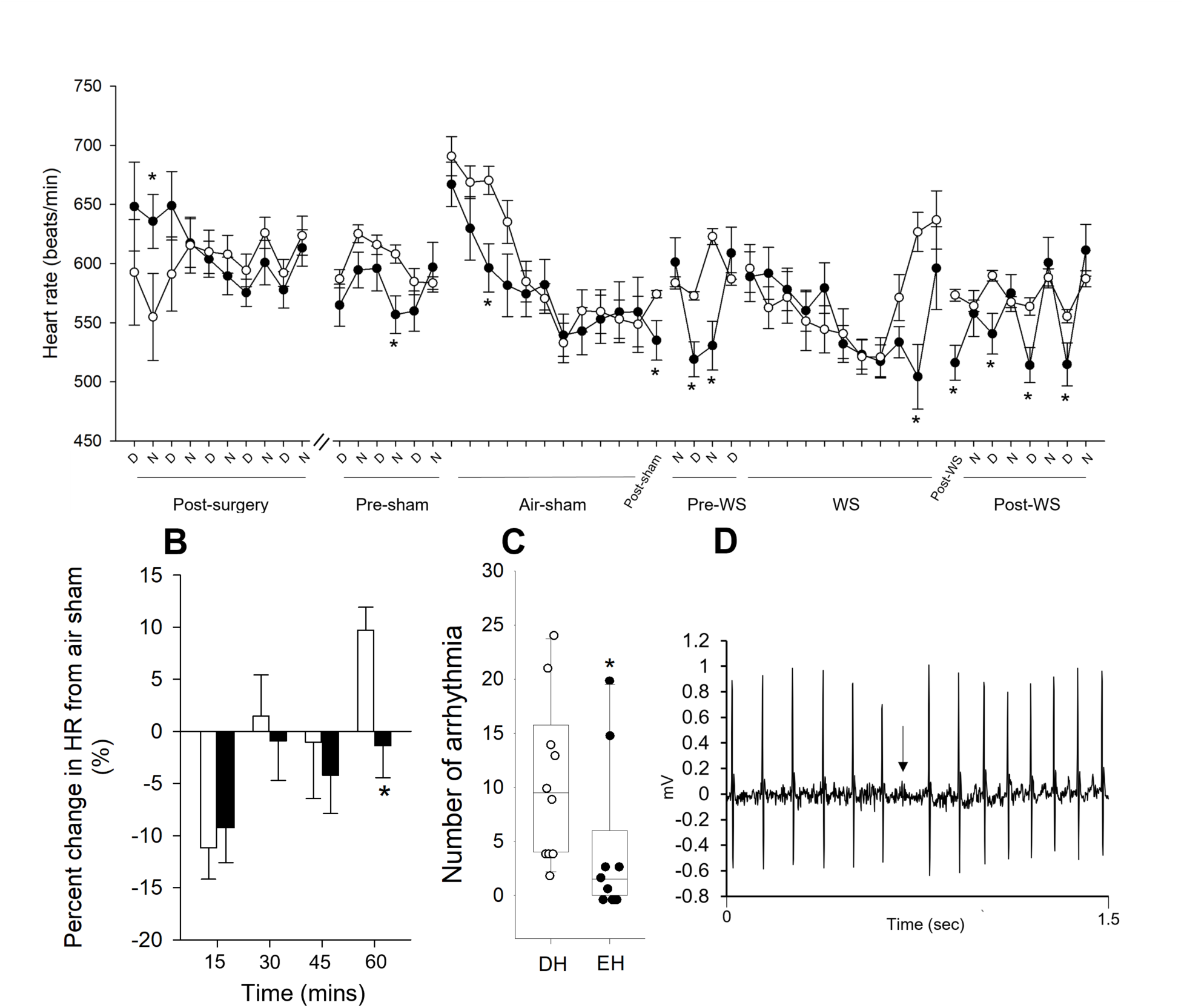
Housing conditions affect day and nighttime heart rate in mice. Although HR was measured continuously after surgery until the end of the study, only data from relevant segments are shown here because the intervening periods had circadian patterns for all animals. **A.** Heart rate was significantly increased during the nighttime in mice housed in an enriched environment (filled circles) immediately after radiotelemetry surgery when compared to depleted housing (open circles). The heart rates of both groups became comparable thereafter. In addition, mice living in an enriched environment had significantly lower heart rates, especially after the air sham and WS exposures, when compared to depleted housing. **B.** During the one-hour exposure to WS, mice living in an enriched housing environment had less change in heart rate relative to the air sham exposure and **C.** significantly fewer non-conducted p-wave or block arrhythmia when compared to depleted mice. **D.** Representative trace showing a non-conducted p-wave. n = 10-12; * Significantly different from DH, p < 0.05.

### Heart rate variability

Table 2 shows the time (SDNN, RMSSD) and frequency (LF/HF) domain HRV for the telemetered animal groups at baseline and during WS exposure. Additional analyses were not performed due to limitations in data acquisition. There were no significant differences between DH and EH except for during the second period of the WS exposure, which corresponded to a trend in increased HR.

Table 2.

**Table 2.**
| Housing | Period | SDNN<br>(msec) | RMSSD<br>(msec) | LF/HF |
| --- | --- | --- | --- | --- |
| Depleted | Baseline | 8.8 ± 1.3 | 7.2 ± 2.0 | 4.1 ± 0.8 |
|  | WS (1<sup>st</sup> 20min) | 13.8 ± 1.0 | 9.3 ± 1.4 | 8.0 ± 1.1 |
|  | WS (2<sup>nd</sup> 20min) | 14.2 ± 0.8 | 8.6 ± 1.1 | 10.2 ± 1.9 |
|  | WS (3<sup>rd</sup> 20min) | 12.9 ± 1.1 | 8.2 ± 1.2 | 7.6 ± 1.2 |
| Enriched | Baseline | 8.9 ± 0.7 | 6.1 ± 1.1 | 4.7 ± 0.9 |
|  | WS (1<sup>st</sup> 20min) | 12.8 ± 1.2 | 8.2 ± 1.1 | 8.3 ± 1.2 |
|  | WS (2<sup>nd</sup> 20min) | 11.3 ± 0.7* | 6.4 ± 0.6 | 9.5 ± 1.6 |
|  | WS (3<sup>rd</sup> 20min) | 13.3 ± 0.9 | 8.3 ± 1.0 | 9.2 ± 1.3 |

\*Significantly different from DH

### Activity

Figure 5 shows the activity of animals, which was blunted for 3-4 days after radiotelemeter surgery. However, it returned to normal thereafter with increased counts/min at night and was reduced during the day. Mice in depleted housing had significantly decreased activity after air sham and WS when compared to animals living in enriched housing. The activity returned to normal with 1 to 2 days of exposure.

**Figure 5.**
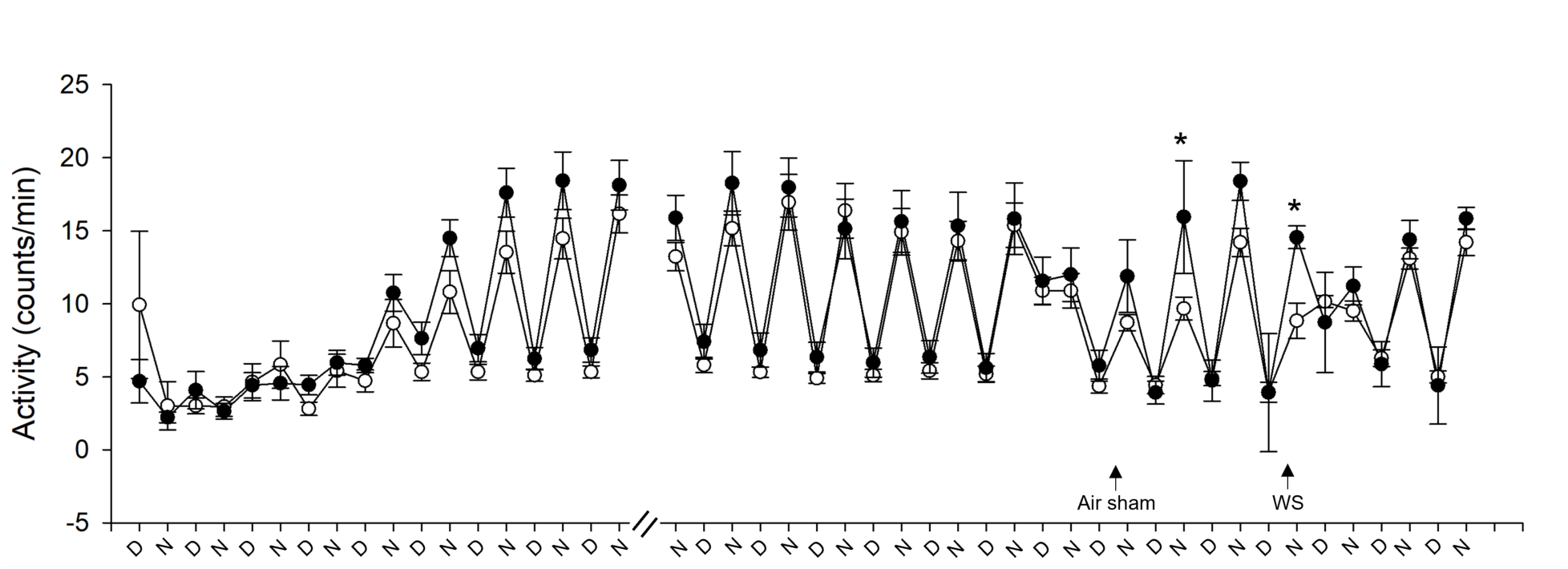
Housing conditions alter activity after air sham and wildfire smoke exposure. All mice experienced blunted activity during the 3 to 4 days after radiotelemeter surgery, but all of them established normal activity thereafter (higher counts/min at night and lower during the day), which continued until the air sham exposure (data shown is not continuous for the entire duration because there was no change). Animals housed in enriched housing (filled circles) had increased activity one day after air sham exposure and immediately after WS when compared to DH (open circles). n = 10-12; * Significantly different from DH, p < 0.05.

### Ventilatory function

Breathing parameters and indicators of airways irritation were measured using whole-body plethysmography immediately after the air sham and the WS exposure in Cohort B. Figure 6 shows the percent change in ventilatory function due to WS relative to air sham. Animals living in enriched housing had significantly less decrease in TV, PIF, and PEF due to WS when compared to animals living in depleted housing. There was also a trend of less increase in Te (p = 0.054).

**Figure 6.**
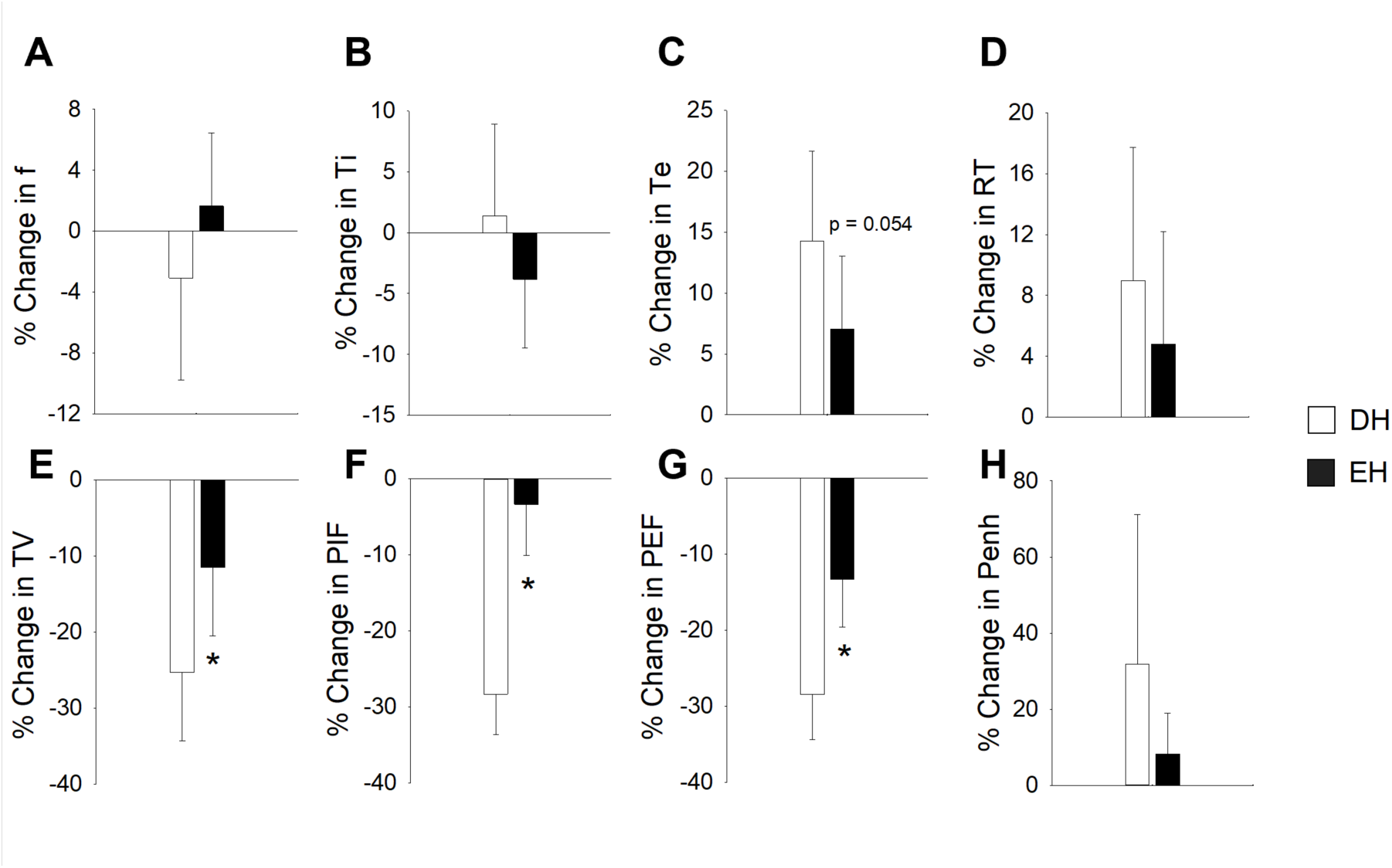
Housing conditions alter the ventilatory response to wildfire smoke exposure. Responses were measured immediately after WS exposure and the percent change from air sham responses were determined. EH mice had significantly less decrease in TV (E), PIF (F), and PEF (G) after WS when compared to DH. There were no other significant differences. n = 8-9; * Significantly different from DH, p < 0.05.

### Gene expression in cardiac tissue

Gene expression was assessed in mouse cardiac tissue from Cohort B using a customized heart gene panel using NanoString technology. Figure 7 shows a volcano plot of differential gene expression in DH and EH mice. The log ratio of the fold change is on the X-axis, and the negative log of p-adj/p-value is on the Y-axis. With a cut-off of 1.25-fold change in expression, seven genes were significantly upregulated (actin alpha2 (ACTA2), androgen receptor (AR), gap junction protein alpha 5 (GJA5), hypocretin receptor 1 (HCRTR1), natriuretic peptide A (NPPA), natriuretic peptide B (NPPB) and P75) and one down-regulated (transient receptor potential cation V1 channel (TRPV1)) in EH mice when compared to DH. When the filter parameter was changed to 1.15-fold change, additional genes were significantly increased due to housing enrichment (voltage-dependent L type calcium channel alpha 1 (CACNA1C), collagen type III alpha 1 (COL3A1), fibronectin 1 (FN1), janus kinase 1 (JAK1), calcium activated potassium channel (KCNN1), nitric oxide synthase 3 (NOS3), superoxide dismutase (SOD), transforming growth factor beta 1 (TGFB1), and transient receptor potential cation M4 channel (TRPM4)). Gene ontology analysis indicated the biological processes and molecular functions that were significantly impacted from the expression changes due to living conditions. These included negative regulation of systemic arterial blood pressure and neuropeptide signaling among others, and hormone activity and binding in the DH group relative to EH. See Table S1 in the Supplemental Material for the full gene expression results.

**Figure 7.**
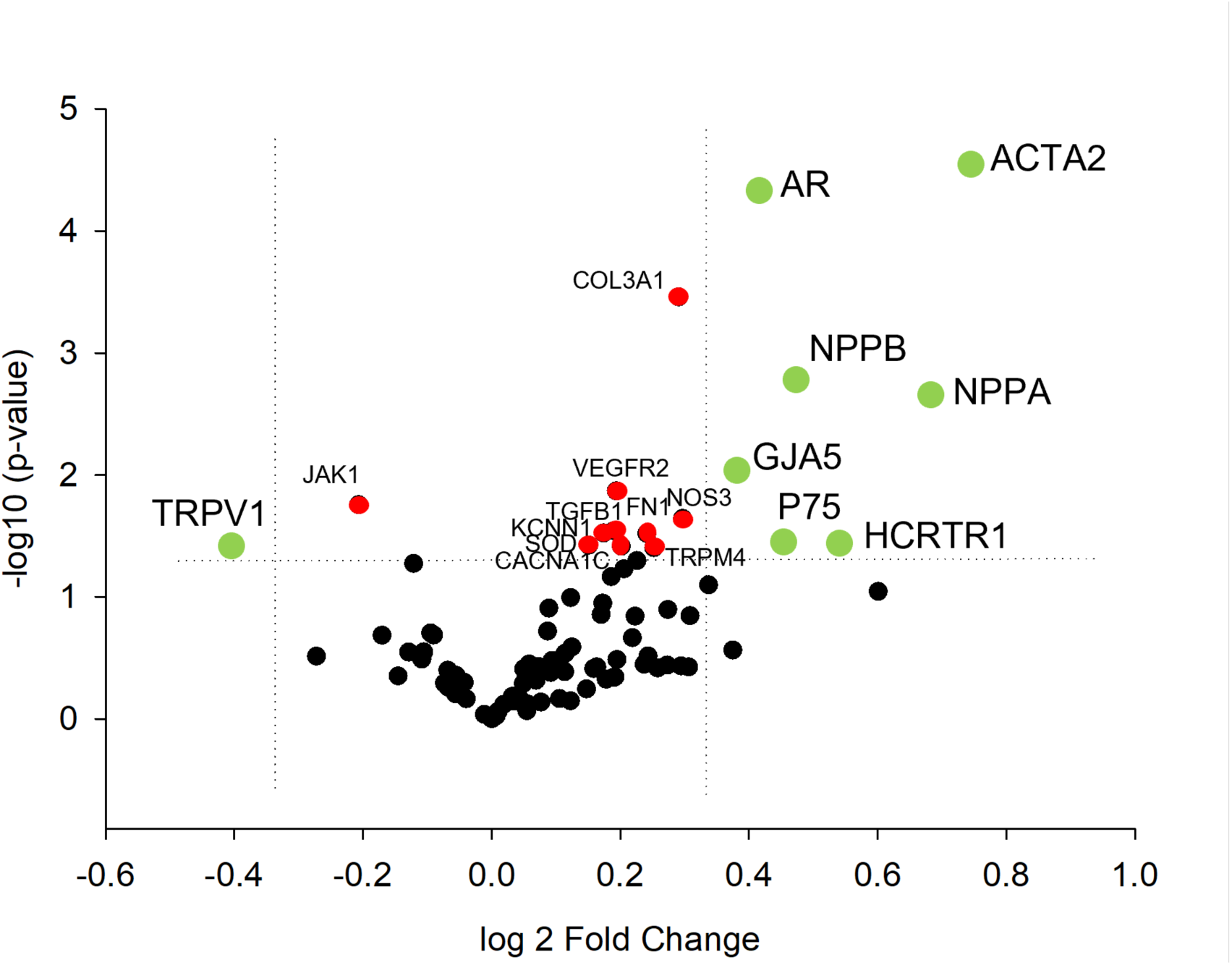
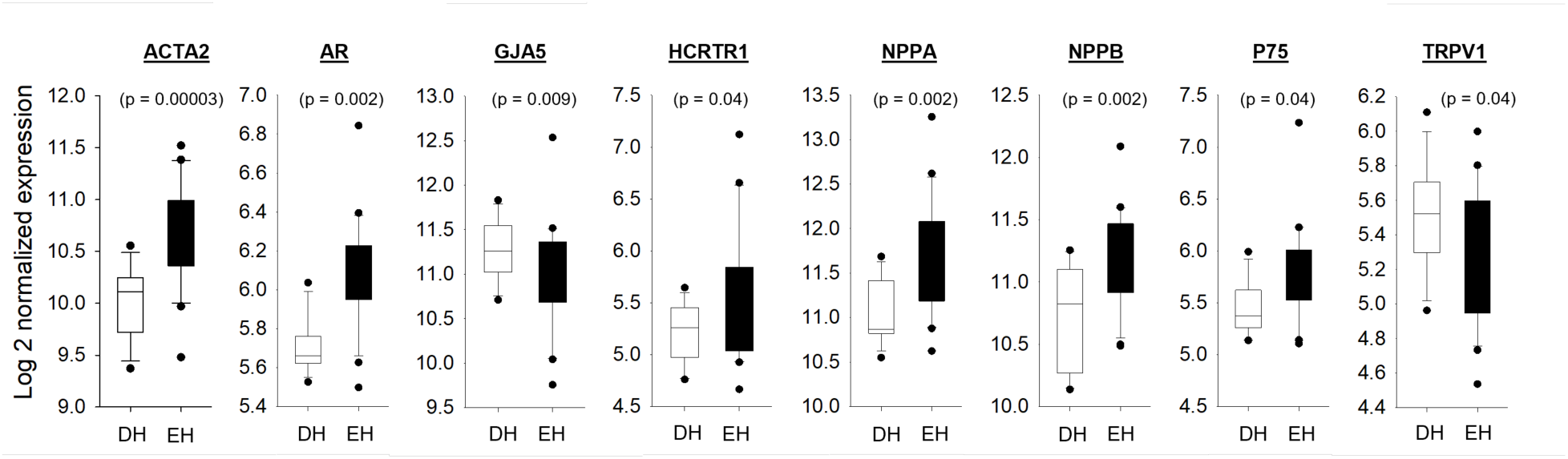
Living conditions alter cardiac gene expression in mice. Volcano plot showing the differences in expression between DH and EH mice. Each dot represents a gene within the comparison performed. The coloring on the dots indicates the clustering information for each gene, those in black are outside the parameters of the filter and not statistically significant. Genes in green were either significantly increased (ACTA, AR, GJA5, HCRTR1, NPPA, NPPB, P75) or decreased (TRPV1) in EH hearts when compared to DH with a 1.25-fold change cut-off. Each are depicted in the box and whisker plot. When the filter was set to 1.15-fold change, additional genes showed significant increases at p < 0.05 (red). n = 11-12; * Significantly different from DH, p < 0.5.

### Bronchoalveolar lavage and serum markers

There were no differences in the number of inflammatory cells or oxidative stress or injury markers in the BAL of any of the groups. CRP was increased in the serum of DH mice when compared to EH and this effect was significantly reduced by WS (See Tables S2 and S3 in Supplemental material, respectively). There were no other differences.

## Discussion

The results of this study clearly show that housing conditions impact overall cardiovascular health and subsequent responses to wildfire-related smoke in mice. Although it has been known for a couple of decades that air pollution exposure increases cardiovascular morbidity and mortality, especially after extreme events like wildfires (Chen et al. 2021; Hadley et al. 2022), the data presented here suggest that daily modifying factors likely modulate this effect. More importantly, living in certain housing conditions for even a short term can alter basic physiology and appears to change the ability of the heart to compensate for various stressors. In addition, our work suggests that depleted housing conditions cause systemic effects, worsen overall body health, and change the behavior of healthy mice, which underscores the importance of where someone lives, not only as a factor of environmental justice, but in the development of remediation strategies.

The place where a person lives has a significant impact on their health and well-being (CDC 2018). Healthy People 2030, which is a federal, has set objectives to improve the health of people across the United States and it includes “Neighborhoods and Built Environment” as a major Social Determinant of Health (SDOH). These determinants contribute to numerous health disparities and inequities and increase the risk of chronic conditions like heart disease, diabetes, and obesity, and more seriously, lower life expectancy (Health People 2030). A longitudinal study showed a higher rate of hypertension due to neighborhood-level socioeconomic deprivation in Dallas (Claudel et al. 2018) while others have shown that dilapidated housing worsens cardiovascular disease due to poor urban outdoor air quality (Shaw 2004; Rajagopalan et al. 2018). Similar impacts of depleted versus enriched housing conditions are documented in mice (Slater and Cao 2015; Bailoo et al. 2018), with the latter improving welfare, overall health, and behavior. Thus, we sought to utilize such a comparative model for the purpose of testing the impacts of air pollution exposure and assessing relative risk of adverse responses.

Although mice housed in enriched conditions had lower initial (before separation in housing) body weights when compared to DH, they had a greater increase in body weight between weeks 2 and 6, and less thereafter. Similar work with the same strain of mice showed that body growth during adolescence is reduced in mice that are individually housed versus those that are socially housed. However, the growth of the individually housed mice increases more significantly during adulthood (Schipper et al. 2020). The data suggest that psychosocial stress and housing conditions play a role in energy balance and consumption. In our mice, the presence of a running wheel likely impacted these endpoints and altered physiological function. Unfortunately, body composition and metabolic assessments were not performed and in the future these endpoints will shed light on the systemic impact of the living environment on the interaction between health and behavior.

Studies in humans indicate that housing conditions are linked to malnutrition and weight gain/obesity (Nobari et al. 2018), yet it is not easy to establish clear correlations and biological plausibility (Bor 2016) in the absence of controlled experiments. Although relatively simplistic, our model showed that depleted housing conditions alter regulation of body temperature and activity level in mice, especially after a stressor. The observations were conducted at temperatures below mouse thermoneutrality (i.e., 30°C), which inherently implies that the animals were expending energy to maintain their core body temperature under those conditions (Gordon 1985). This response would likely be strained in the absence of environmental enrichment, which has been shown to alleviate thermal discomfort (Gaskill et al. 2013), and/or the presence of a stressor. In the case of the latter, one study showed that mice exposed to cigarette smoke had pronounced hypothermia (van Eijl et al. 2006), possibly due to the restraint involved, while another demonstrated similar temperature dysregulation because of social isolation (Kalliokoski et al. 2014). Thus, post-surgical hypothermia in our DH mice likely pointed to not only the impacts of stress, but also the benefits of housing enrichment for the EH group. Moreover, EH mice returned to normal activity levels after both the air sham and smoke exposures, suggesting better resiliency and coping when it comes to stress.

Today, there is sufficient evidence implicating deteriorated or depleted housing in the development of cardiovascular diseases (Sims et al. 2020). Similar to changes in activity and body temperature, heart rate was significantly lower after telemetry surgery in mice housed in depleted conditions when compared to enriched. This is likely part of the mouse hypothermic response (Gordon 2017), which in other models of stress can be ameliorated with thermoneutrality (Koizumi et al. 1992), and also points to impaired resiliency and perhaps increased risk. Heart rate in DH mice did, however, return to levels on par with EH animals after recovery from surgery. On the other hand, DH animals had elevated heart rates during air sham, immediately after air sham, during WS, and after WS. Persistently elevated heart rate is a risk factor for adverse cardiovascular events (Palatini and Julius 2004) and it is independently associated with morbidity and mortality, even in healthy individuals (Hjalmarson 2007).

Thus, here we show that enrichment lessens cardiovascular responses, potentially through regulation of the autonomic nervous system and maintenance of resting physiology. It is worth noting that elevated heart rate during and after stressful events is particularly important when predicting the development of future disease (Palatini 2011). A recent study showed that life events and psychosocial stress are associated with elevated heart rate (Schneider et al. 2021). Furthermore, lower socioeconomic (Zhang et al. 2016) and housing status (Sims et al. 2020) contribute to elevated resting heart rate and poorer overall health. Similar effects have been reported in rodents due to housing inadequacy and stressful challenges (Spani et al. 2002; Sharp et al. 2014). Although, we exposed mice to a fairly high concentration of eucalyptus wildfire smoke as a stressor, we are the first to show that not only does heart rate remain elevated post-exposure due to depleted housing, but that it also increases the incidence of cardiac arrhythmia.

In people, greenspace has been recognized as a positive contributor to well-being, and a number of studies have shown that this effect is due to improvements in heart rate variability, which reflects autonomic balance in the heart (Lee et al. 2014; de Brito et al. 2020). Unfortunately, it is unclear whether significant changes in HRV contributed to the effects observed in our study. Models of housing enrichment in rodents suggest the effect on HRV is gradual, consistent with findings indicating autonomic function becomes altered by enrichment over the long term (Brauner et al. 2010). In any case, the higher HRV in DH mice during WS exposure may be due to airways irritation and the activation of autonomic reflexes, which we have previously shown in mice (Kurhanewicz et al 2018). In rats, exposure to noise stress and then ozone also resulted in an increase in HRV (Hazari et al. 2021).

These responses, at least in terms of what we observed in the current study, may be driven by changes in the expression and function of certain receptors and channels that mediate chronotropy and inotropy in the heart, especially over the long term. We found that EH led to the upregulation of natriuretic peptide A (NPPA) and B (NPPB), which have been shown to limit cardiomyocyte hypertrophy and death, and reduce cardiac fibrosis (Forte et al. 2022), hypocretin receptor 1 (HCRTR1), the deficiency in which leads to poorer cardiac function (Perez et al. 2015), and other genes that regulate cardiovascular function. On the other hand, transient receptor potential cation channel V1 (TRPV1), which plays a role in the progression of cardiac hypertrophy (Buckley and Stokes 2011), temperature related anxiety (Lima et al. 2022), and general stress (Ho et al. 2012) was down-regulated. Furthermore, TRPM4, which is necessary for regulation of positive inotropy and blood pressure (Mathar et al. 2010, 2013), was also upregulated. These finding suggest that enriched housing elicits a gene expression profile that may not only be protective of the heart, but one that also reduces stress responsiveness. Similar patterns of change were observed in the Multi-Ethnic Study of Atherosclerosis (MESA), which showed that neighborhood socioeconomic disadvantage leads to increased stress and inflammation gene expression (Smith et al. 2017).

Interestingly, TRP channels are also known to mediate the irritant response of the airways to inhaled pollutants. Thus, it was no surprise that we observed reduced ventilatory responses, especially in peak airflow, in mice housed in enriched conditions. Although it is expected that a reduction in ventilation is protective during an inhaled exposure to air pollution, here it is unlikely that EH animals have impaired airway defensive reflexes, as indicated by the fact that there was no difference in Penh. Instead, it may be that EH mice are able to better compensate for the irritant exposure than DH animals.

In conclusion, the results of this study show that living conditions impact overall cardiovascular health in mice and impact the response to air pollution exposure. As demonstrated here, enriched housing appears to change the body’s ability to compensate for stress and returns function to normal more rapidly than if the conditions are depleted. This paradigm does not show that housing has a significant impact on general markers of injury and oxidative stress or lung inflammation following a robust wildfire smoke exposure. However, the molecular changes in the heart suggest that long-term effects would impact responses to environmental exposures. Although further work is needed to characterize the systemic changes, these data point to the importance of non-chemical factors in determining health and future risk.

