## Supplementary Materials for "The effect of enriched versus depleted housing on eucalyptus smoke-induced cardiovascular dysfunction in mice"

Table S1. Normalized fold-differences in cardiac gene expression in EH group relative to DH group. P-values are shown unadjusted (p value) and following statistical adjustment for false discovery rate (p adj).

| *Gene* | *Description* | *Fold Change* | *log2-Fold Change* | *p value* | *p adj* |
|---|---|---|---|---|---|
| ACE | Angiotensin I converting enzyme | 1.08218 | 0.113936 | 0.291593 | 0.613328 |
| ACTA2 | Actin $\alpha$2_ smooth muscle | 1.67624 | 0.74523 | 2.96E-05 | 0.002161 |
| ACTC1 | Actin $\alpha$1_ cardiac muscle | 1.15341 | 0.205904 | 0.059036 | 0.255822 |
| ADRA1A | Adrenoceptor $\alpha$ 1A | -1.00796 | -0.01144 | 0.922002 | 0.953434 |
| ADRA1B | Adrenoceptor $\alpha$ 1B | 1.02547 | 0.036286 | 0.71966 | 0.798647 |
| ADRA1D | Adrenoceptor $\alpha$ 1D | 1.164 | 0.219088 | 0.216993 | 0.54851 |
| ADRB1 | Adrenoceptor $\beta$ 1 | 1.00488 | 0.007016 | 0.9453 | 0.962011 |
| ADRB2 | Adrenoceptor $\beta$ 2 | 1.07585 | 0.105476 | 0.683819 | 0.798142 |
| ADRB3 | Adrenoceptor V 3 | 1.22648 | 0.294525 | 0.369532 | 0.613328 |
| AGTR1B | Angiotensin II receptor type 1 | 1.03866 | 0.054717 | 0.86117 | 0.909771 |
| AHR | Aryl hydrocarbon receptor | -1.04799 | -0.06763 | 0.553733 | 0.699858 |
| AR | Androgen receptor | 1.33365 | 0.415379 | 4.75E-05 | 0.002161 |
| ATP2A | ATPase $Ca^{2+}$ transporting_ cardiac muscle_ fast twitch 1 | 1.23665 | 0.306443 | 0.376612 | 0.613328 |
| ATP5A1 | ATP synthase $H^{+}$ transporting_ mitochondrial F1 complex $\alpha$1 | -1.06797 | -0.09487 | 0.198505 | 0.537708 |
| CACNA1C | Calcium channel voltage-dependent L type $\alpha$ 1C subunit | 1.14998 | 0.201604 | 0.038542 | 0.2001 |
| CAT | Catalase | 1.06356 | 0.088904 | 0.123912 | 0.414345 |
| CCL2 | Chemokine (C-C motif) ligand 2 | 1.1313 | 0.177977 | 0.476219 | 0.646806 |
| CCL3 | Chemokine (C-C motif) ligand 3 | 1.08857 | 0.122436 | 0.713257 | 0.798647 |
| COL11A1 | Collagen type XI $\alpha$1 | 1.0 | 0 | 1 | 1 |
| COL3A1 | Collagen type III $\alpha$1 | 1.22349 | 0.291006 | 0.000348 | 0.010556 |
| CXCL8 | Chemokine (C-X-C motif) ligand 8 | 1.23866 | 0.308775 | 0.143046 | 0.424475 |
| DMD | Dystrophin | 1.12721 | 0.172757 | 0.113231 | 0.396309 |
| EDN1 | Endothelin 1 | 1.03962 | 0.056051 | 0.761646 | 0.816544 |
| F2R | Coagulation factor II (thrombin) receptor | 1.05099 | 0.071746 | 0.374267 | 0.613328 |

| | | | | | |
|---|---|---|---|---|---|
| FABP3 | Fatty acid binding protein 3 | -1.09362 | -0.1291 | 0.28452 | 0.613328 |
| FN1 | Fibronectin 1 | 1.1822 | 0.241475 | 0.030341 | 0.2001 |
| FOS | FBJ murine osteosarcoma viral oncogene homolog | 1.29715 | 0.375344 | 0.273437 | 0.613328 |
| GJA1 | Gap junction protein $\alpha$1 | -1.10604 | -0.1454 | 0.44753 | 0.635185 |
| GJA5 | Gap junction protein $\alpha$5 | 1.30198 | 0.380705 | 0.009239 | 0.140129 |
| GPX1 | Glutathione peroxidase 1 | -1.00217 | -0.00312 | 0.962011 | 0.962011 |
| GSTP1 | Glutathione S-transferase pi 1 | 1.04108 | 0.058082 | 0.356389 | 0.613328 |
| HCRT | Hypocretin (orexin) neuropeptide precursor | 1.0 | 0 | 1 | 1 |
| HCRTR1 | Hypocretin (orexin) receptor 1 | 1.45421 | 0.540238 | 0.037902 | 0.2001 |
| HCRTR2 | Hypocretin (orexin) receptor 2 | 1.0 | 0 | 1 | 1 |
| HMOX1 | Heme oxygenase 1 | 1.06981 | 0.097348 | 0.361876 | 0.613328 |
| HSPA1A | Heat shock protein 1A | 1.05447 | 0.076522 | 0.731832 | 0.80237 |
| ICAM1 | Intercellular adhesion molecule 1 | 1.17887 | 0.237407 | 0.357408 | 0.613328 |
| IL-1ALPHA | Interleukin 1$\alpha$ | 1.11609 | 0.158456 | 0.388624 | 0.613328 |
| IL-1BETA | Interleukin 1$\beta$ | 1.14442 | 0.194612 | 0.3272 | 0.613328 |
| IL-6 | Interleukin 6 | 1.19601 | 0.258233 | 0.384518 | 0.613328 |
| JAK1 | Janus kinase 1 | -1.15416 | -0.20684 | 0.017547 | 0.1996 |
| KCNH2 | Potassium voltage gated channel subfamily H member 2 | 1.06572 | 0.091829 | 0.417643 | 0.623041 |
| KCNJ12 | Potassium inwardly rectifying channel subfamily J member 12 | -1.03964 | -0.05608 | 0.625117 | 0.768724 |
| KCNN1 | Potassium channel calcium activated intermediate/small conductance subfamily N$\alpha$ member 1 | 1.12814 | 0.17394 | 0.030046 | 0.2001 |
| KCNQ1 | Potassium voltage gated channel KQT-like subfamily Q member 1 | 1.12505 | 0.169991 | 0.140085 | 0.424475 |
| LDHA | Lactate dehydrogenase A | 1.16959 | 0.225998 | 0.050567 | 0.24219 |
| MAOA | Monoamine oxidase A | 1.03029 | 0.043048 | 0.682609 | 0.798142 |
| MAOB | Monoamine oxidase B | 1.00692 | 0.009948 | 0.869781 | 0.909771 |
| MAPK1 | Mitogen-activated protein kinase 1 | -1.03004 | -0.04269 | 0.503584 | 0.658027 |
| MAPK8 | Mitogen-activated protein kinase 8 | 1.03499 | 0.049617 | 0.390912 | 0.613328 |
| MMP1 | Matrix metallopeptidase 1 | 1.13999 | 0.189016 | 0.460683 | 0.635185 |
| MMP9 | Matrix metallopeptidase 9 | 1.10758 | 0.147416 | 0.570595 | 0.71129 |

| | | | | | |
|---|---|---|---|---|---|
| MT1X | Metallothionein 1X | -1.05213 | -0.07331 | 0.512483 | 0.658027 |
| MYC | v-myc avian myelocytomatosis viral oncogene homolog | 1.20932 | 0.274195 | 0.127491 | 0.414345 |
| MYH6 | Myosin heavy chain 6 | 1.03465 | 0.049138 | 0.513406 | 0.658027 |
| MYH7 | Myosin heavy chain 7 | 1.26357 | 0.337502 | 0.079933 | 0.316255 |
| MYL2 | Myosin light chain 2 | -1.03919 | -0.05546 | 0.441185 | 0.635185 |
| MYL3 | Myosin light chain 3 | 1.02069 | 0.029538 | 0.701663 | 0.798142 |
| NFKB1 | Nuclear factor kappa B | 1.02255 | 0.086946 | 0.191304 | 0.537708 |
| NGF | Nerve growth factor | 1.08188 | 0.032169 | 0.655429 | 0.795254 |
| NOS2 | Nitric oxide synthase 2 | 1.11986 | 0.113547 | 0.412209 | 0.623041 |
| NOS3 | Nitric oxide synthase 3 | 1.22877 | 0.163323 | 0.37619 | 0.613328 |
| NPPA | Natriuretic peptide A | 1.60321 | 0.297211 | 0.022794 | 0.2001 |
| NPPB | Natriuretic peptide B | 1.38844 | 0.680959 | 0.002264 | 0.0412 |
| NPY | Neuropeptide Y | 1.0 | 0.473462 | 0.00167 | 0.037989 |
| NQO1 | NAD(P)H dehydrogenase quinone 1 | 1.06721 | 0 | 1 | 1 |
| NR3C1 | Nuclear receptor subfamily 3 group C1 (glucocorticoid receptor) | -1.08784 | 0.093839 | 0.334154 | 0.613328 |
| NR3C2 | Nuclear receptor subfamily 3 group C2 | 1.02963 | -0.12147 | 0.053528 | 0.243554 |
| P75 | P75 | 1.36966 | 0.042124 | 0.701002 | 0.798142 |
| PDE3A | Phosphodiesterase 3A | 1.08883 | 0.453819 | 0.035032 | 0.2001 |
| PECAM1 | Platelet/endothelial cell adhesion molecule 1 | 1.04885 | 0.122776 | 0.10195 | 0.371098 |
| RYR1 | Ryanodine receptor 1 (skeletal) | 1.0 | 0.068806 | 0.485554 | 0.649785 |
| RYR2 | Ryanodine receptor 2 (cardiac) | 1.00407 | 0 | 1 | 1 |
| SCN3B | Sodium channel voltage gated type III beta subunit | 1.20836 | 0.00586 | 0.954529 | 0.962011 |
| SCN5A | Sodium channel voltage gated type V alpha subunit | -1.02779 | 0.273047 | 0.36375 | 0.613328 |
| SOD | Superoxide dismutase | 1.11013 | -0.03954 | 0.688237 | 0.798142 |
| STAT1 | Signal transducer and activator of transcription 1 | 1.01302 | 0.15073 | 0.037895 | 0.2001 |
| TAC1 | Tachykinin precursor 1 | 1.0 | 0.018656 | 0.762706 | 0.816544 |
| TACR1 | Tachykinin receptor 1 | 1.14188 | 0 | 1 | 1 |
| TGFB1 | Transforming growth factor beta 1 | 1.14062 | 0.191416 | 0.456087 | 0.635185 |

| | | | | | |
|---|---|---|---|---|---|
| THBS1 | Thrombospondin 1 | 1.51684 | 0.189812 | 0.028788 | 0.2001 |
| TNF-ALPHA | Tumor necrosis factor | 1.18384 | 0.601071 | 0.090591 | 0.343491 |
| TNNC1 | Troponin C type 1 (slow) | 1.04237 | 0.243469 | 0.306305 | 0.613328 |
| TNNI3 | Troponin I type 3 (cardiac) | -1.06471 | 0.059861 | 0.439319 | 0.635185 |
| TNNT2 | Troponin T type 2 (cardiac) | -1.04847 | -0.09046 | 0.206357 | 0.537708 |
| TPM1 | Tropomyosin 1 (alpha) | -1.07746 | -0.06829 | 0.401236 | 0.618856 |
| NTRKA | Neurotrophic tyrosine kinase receptor type 1 | 1.0 | -0.10764 | 0.297877 | 0.613328 |
| TRPA1 | Transient receptor potential cation channel A1 | -1.20779 | 0 | 1 | 1 |
| TRPC6 | Transient receptor potential cation channel C6 | 1.0 | -0.27237 | 0.307342 | 0.613328 |
| TRPM4 | Transient receptor potential cation channel M4 | 1.19118 | 0 | 1 | 1 |
| TRPV1 | Transient receptor potential cation channel V1 | -1.32459 | 0.252395 | 0.03958 | 0.2001 |
| TXN | Thioredoxin | -1.07824 | -0.40554 | 0.038112 | 0.2001 |
| UBB | Ubiquitin B | -1.07599 | -0.10867 | 0.32475 | 0.613328 |
| VCAM1 | Vascular cell adhesion molecule 1 | 1.16721 | -0.10566 | 0.283129 | 0.613328 |
| VEGFA | Vascular endothelial growth factor A | 1.09022 | 0.223065 | 0.144601 | 0.424475 |
| VEGFB | Vascular endothelial growth factor B | -1.12535 | 0.124613 | 0.258881 | 0.613328 |
| KDR | Kinase insert domain receptor | 1.14371 | -0.17038 | 0.206811 | 0.537708 |
| ZYX | Zyxin | 1.13739 | 0.193717 | 0.013542 | 0.176049 |

### Table S2. Bronchoalveolar lavage biomarkers

| | Bronchoalveolar lavage | | | |
|---|---|---|---|---|
| | LDH | NAG | SOD | Prot |
| DH-FA | 33.8 ± 15.1 | 8.6 ± 0.2 | 0.23 ± 0.05 | 51.2 ± 11.1 |
| DH-WS | 32.7 ± 10.6 | 8.8 ± 0.1 | 0.51 ± 0.12 | 73.7 ± 18.6 |
| EH-FA | 42.1 ± 13.4 | 8.6 ± 0.1 | 0.54 ± 0.19 | 56.2 ± 13.4 |
| EH-WS | 68.1 ± 17.7 | 8.6 ± 0.2 | 0.55 ± 0.17 | 49.6 ± 9.7 |

LDH = lactate dehydrogenase (U/l)
NAG = N-acetyl-beta-D-glucosaminidase (U/l)
SOD = superoxide dismutase (U/ml)
Prot = protein (ug/ml)

### Table S3. Serum and bronchoalveolar lavage biomarkers

| | Serum | | | | | | |
|---|---|---|---|---|---|---|---|
| | Chol | CK | CRP | GPX | GTR | HDL | LDH |
| DH-FA | 99.8 ± 8.1 | 331 ± 99 | 0.6 ± 0.03 | 0.5 ± 0.03 | 0.04 ± 0.008 | 37.7 ± 2.0 | 141 ± 67 |
| DH-WS | 96.4 ± 6.6 | 458 ± 300 | 0.07 ± 0.02 | 0.6 ± 0.1 | 0.03 ± 0.005 | 36.1 ± 2.1 | 310 ± 125 |
| EH-FA | 96.1 ± 8.7 | 689 ± 264 | 0.09 ± 0.02* | 0.5 ± 0.02 | 0.03 ± 0.006 | 34.7 ± 2.8 | 228 ± 68 |
| EH-WS | 80.6 ± 7.1 | 171 ± 87 | 0.08 ± 0.02 | 0.5 ± 0.03 | 0.03 ± 0.007 | 36.8 ± 1.8 | 265 ± 59 |

Chol = cholesterol (mg/dl)
CK = creatine kinase (U/l)
CRP = C-reactive protein
GPX = glutathione peroxidase (IU/ml)
GTR = glutathione reductase (IU/ml)
HDL = high-density lipoprotein (mg/dl)
LDH = lactate dehydrogenase (U/l)
